# Quantification of bacterial DNA in blood using droplet digital PCR: a pilot study

**DOI:** 10.1101/2022.12.02.518639

**Authors:** Ana P. Tedim, Irene Merino, Alicia Ortega, Marta Domínguez-Gil, José Maria Eiros, Jesús F. Bermejo-Martín

**Affiliations:** Group for Biomedical Research in Sepsis (BioSepsis). Instituto de Investigación Biomédica de Salamanca, (IBSAL), Paseo de San Vicente, 58-182, 37007 Salamanca, Spain; Hospital Universitario Río Hortega, Calle Dulzaina, 2, 47012 Valladolid, Spain; Microbiology Department, Hospital Universitario Río Hortega, Calle Dulzaina, 2, 47012 Valladolid, Spain; Centro de Investigación Biomédica en Red en Enfermedades Respiratorias (CiberES), CB22/06/00035, Instituto de Salud Carlos III, Av. de Monforte de Lemos, 3-5, 28029 Madrid, Spain

**Keywords:** Bacterial DNA quantification, Bloodstream infections, Droplet digital PCR, ddPCR

## Abstract

**Aim:** To use genus/species-specific genes droplet digital PCR (ddPCR) assays to detect/quantify bacterial DNA from *Escherichia coli*, *Klebsiella pneumoniae*, *Staphylococcus aureus* and *Enterococcus* spp in blood samples.

**Methods and Results:** Bacterial DNA from clinical strains (4<n<12) was extracted, quantified and diluted (10-0.0001ng/μL) and ddPCR assays were performed in triplicate. These ddPCR assays showed low replication variability, low detection limit (1–0.1pg/μL) and high genus/species specificity. ddPCR assays were also used to quantify bacterial DNA obtained from spiked blood (1×104-1CFU/mL) of each bacterial genus/species. Comparison between ddPCR assays and bacterial culture was performed by Pearson correlation. There was an almost perfect correlation (r≥0.997, p≤0.001) between the number of CFU/mL from bacterial culture and the number of gene copies/mL detected by ddPCR. The time from sample preparation to results was determined to be 3.5-4h.

**Conclusions:** The results demonstrated the quantification capacity and specificity of the ddPCR assays to detect/quantify four of the most important bloodstream infection (BSI) bacterial pathogens directly from blood.

**Significance and Impact:** This pilot study results reinforce the potential of ddPCR for the diagnosis and/or severity stratification of BSI. Applied to patients’ blood samples it can improve diagnosis and diminish sample-to-results time, improving patient care.

## Introduction

Bloodstream infections (BSIs) caused by bacteria associated to sepsis are among the leading causes of mortality, particularly in critically ill patients ^1,2^. The gold standard method for the microbiological diagnosis of BSIs is still blood culture, which is slow, cannot detect viruses, and only yields positive results in one-third of suspected BSIs and sepsis cases ^1,3,4^.

Due to the high mortality of BSIs and sepsis, early administration of empiric broad-spectrum antimicrobials is normally the first step in the treatment of these infections, but in many occasions this it is not accurate for the actual bacterial pathogen causing BSI or sepsis, leading to therapeutic failure and death ^1–3,5^. Therefore, it is imperative to identify, in a timely manner, the infectious agent causing BSI or infection leading to sepsis (bacteria, fungi, or virus) as well as possible associated antimicrobial resistances ^2,6^, in order to administer tailored antimicrobial therapy. Thus, it is necessary to develop rapid, sensitive, and accurate molecular diagnostic methods to identify and quantify pathogens and their antimicrobial resistances directly in the blood ^2–4^.

Furthermore, time to blood culture positivity has been suggested as a surrogate marker of blood bacterial load and associated with poor clinical outcome in BSIs ^7,8^. Therefore, the early identification and quantification of bacterial DNA load in blood of patients with suspected BSIs or sepsis could help to assess illness severity and further guide patient’s treatment ^2,6,9^.

Droplet digital PCR (ddPCR) is a next-generation PCR method, with great precision and accuracy, that allows absolute quantification of target gene(s) without a standard curve and little interference from normal PCR inhibitors ^6^. These characteristics make ddPCR an ideal method for the detection and quantification of pathogens directly from blood or other clinical samples ^2,6^ in patients with suspected BSI and sepsis. Nevertheless, very few studies have validated its performance to assist in BSI diagnosis ^6^.

The aim of this work was to detect and quantify bacterial DNA from four of the most common BSIs pathogens (*Escherichia coli, Klebsiella pneumoniae, Staphylococcus aureus* and *Enterococcus* spp) directly from blood using genus/species specific genes ddPCR assays.

## Material and methods

### Bacterial strains and DNA extraction

Clinical strains, identified as belonging to the bacterial species of interest by VITEK^®^ MS (Biomérieux, Marcy-l’Étoile, France), were recovered from de Microbiology Department of Hospital Universitario Río Hortega, Spain, from May 2019 and February 2020 (Table 1). No personal data from patients were recorded, and strains were anonymized on collection. Strains were plated on Columbia blood agar (Oxoid, Basingtoke, Hampshire, United Kingdom) and colonies picked and stored at −80°C. This article does not contain any studies with human or animal participants performed by any of the authors.

**Table 1.** Bacterial species and number of clinical strains used in the study.

| Species | No of clinical strains |
| --- | --- |
| *Escherichia coli* | **10** |
| *Klebsiella pneumoniae* | **12** |
| *Citrobacter braakii* | 1 |
| *Enterobacter cloacae* | 1 |
| *Klebsiella oxytoca* | 3 |
| *Morganella morganii* | 3 |
| *Proteus mirabilis* | 3 |
| *Staphylococcus aureus* | **10** |
| *Staphylococcus constellatus* | 1 |
| *Staphylococcus epidermidis* | 4 |
| *Staphylococcus haemolyticus* | 2 |
| *Staphylococcus hominis* | 2 |
| *Staphylococcus simulans* | 2 |
| *Staphylococcus saprophyticus* | 1 |
| *Enterococcus faecalis* | **6** |
| *Enterococcus faecium* | **4** |
| *Streptococcus agalactiae* | 8 |

Strains were plated on Columbia blood agar (Oxoid, Basingtoke, Hampshire, United Kingdom) and genomic DNA from bacterial colonies was obtained using the Promega Wizard Genomic DNA Purification Kit (Promega, Madison WI, USA). DNA concentrations were measured with Nanodrop (Thermo Scientific, Waltham MA, USA). Serial dilutions were performed with RNase free water to obtain working bacterial DNA concentrations of 10, 1, 0.1, 0.01, 0.001 and 0.0001 ng/μL.

### Description of ddPCR assays

Primers and probes used for ddPCR assays are listed in Table 2. All primers and probes (Integrated DNA Technologies, Coralville IA, USA) have previously been used to detect *E. coli, K pneumoniae, S. aureus* and *Enterococcus* spp. with either real-time PCR or ddPCR ^10–12^. Probes for Gram-negative bacteria (*E. coli*, and *K pneumoniae*) were labelled with HEX fluorophore and probes for Gram-positive bacteria (*S. aureus*, and *Enterococcus* spp.) were labelled with FAM fluorophore.

**Table 2.**
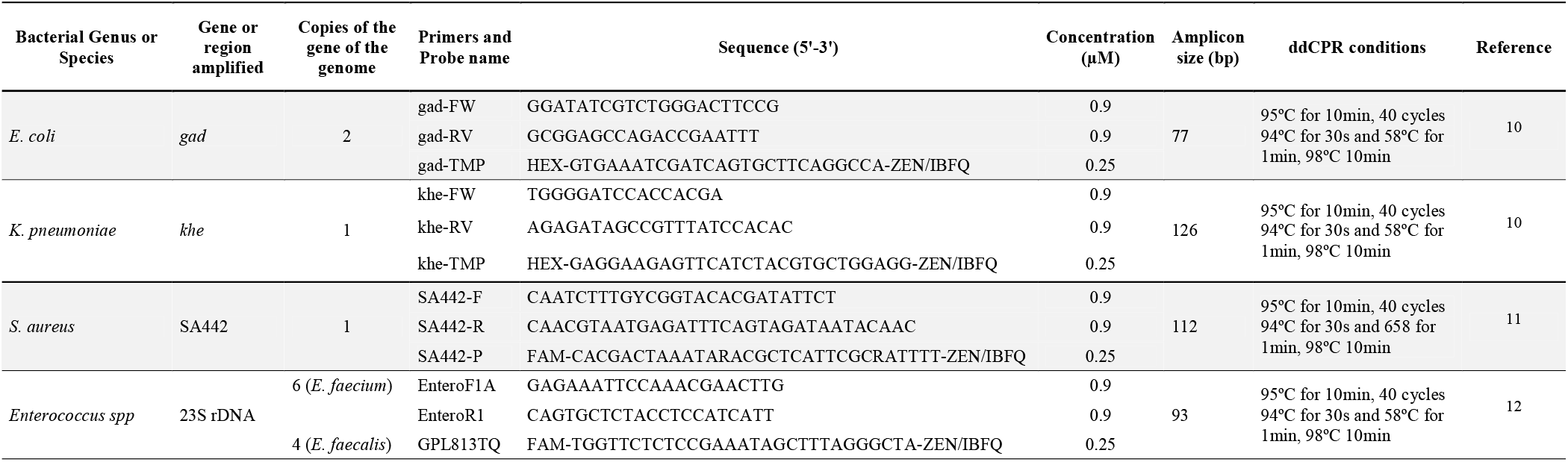
Primers and Probes used in the study.

All the assays were performed using the QX200 Droplet Digital PCR system (Bio-Rad Laboratories Inc., Pleasanton, CA, USA) according to manufacturer instructions. Briefly, template DNA (2.5μL) was added to 18.5μL of mastermix containing ddPCR Supermix for probes no dUPTs (1x), forward and reverse primer (0.9μM each) and probe (0.25μM). This mastermix was placed into a QX200 droplet generator in order to generate droplets. The generated droplet emulsion was transferred to a new 96-well PCR plate and amplified in a C1000 Touch Thermal Cycler (Bio-Rad Laboratories Inc., Pleasanton, CA, USA). The amplification protocol was as follows: 95°C for 10 minutes, 40 cycles at 94°C for 30s and 56-62°C (gradient) for 1 minute, and a final cycle at 98°C for 10 minutes. After gene amplification, the plate was transferred to and read in a QX200 droplet reader (Bio-Rad Laboratories Inc., Pleasanton, CA, USA).

In order to evaluate the performance of primers and probes in the detection of the above-mentioned bacterial species, developmental ddPCR assays were first performed as simplex ddPCR and then *E. coli* and *S. aureus* were tested as duplex ddPCR and *K. pneumoniae* and *Enterococcus (Enterococcus faecium* and *Enterococcus faecalis*), using one clinical strain. In the developmental assays different DNA concentrations (10, 1, 0.1, 0.01, 0.001 and 0.0001 ng/μL) were tested in triplicate.

After determining the ideal concentration to use, further clinical strains (Table 1) were used to evaluate the precision of the test in detecting the target bacteria in the clinical setting and against related isolates. In order to evaluate primers specificity other bacteria similar to *E. coli* and *K. pneumoniae, S. aureus* and *Enterococcus* spp. were included (Table 1) in the assay.

### Spiked blood ddPCR assays

One clinical strain for *E. coli, K. pneumoniae, S. aureus, E. faecium* and *E. faecalis* was inoculated in blood agar plate and incubated overnight at 37°C. One colony for each bacterial species was inoculated in thioglycolate broth (Biomérieux, Marcy-l’Étoile, France) and incubated overnight at 37°C. Each bacterial culture was serial dilute (1:10) in saline 0.9% (1×10^8^ to 1 CFU/mL), and 20μL were plated in duplicate in blood agar plates and incubated overnight at 37°C. Colonies were counted to calculate the CFU/mL of the initial thioglycolate broth culture. Blood was spiked with saline dilutions of 1×10^5^, 1×10^4^, 1×10^3^, 1×10^2^ and 10 CFU/mL in order to obtain blood spiked with 1×10^4^, 1×10^3^, 1×10^2^, 10 and 1 CFU/mL. DNA was extracted from spiked blood (1×10^4^ to 1 CFU/mL) using MolYsis Basic5 DNA extraction kit (Molzym, Bremen, Germany) combined with QIAamp UCP Pathogen Mini Kit (Qiagen, Venlo, Netherlands) following manufacturer instructions. For ddPCR assays 2.5μL of each DNA extracted from spiked blood (1×10^4^ to 1 CFU/mL) were tested in the previously described ddPCR duplex assays (see above). The number of copies of bacterial housekeeping genes present for each bacterial genome were calculated taking into account the number of copies each bacterial gene has in the species genome (Table 2). These experiments were performed in duplicate.

### Statistics

Pearson correlation coefficient was used to compare the CFU/mL results obtained from bacterial cultures with the average of copy number/mL obtained from the two spiked blood experiments.

## Results

The best annealing temperature for all the simplex assays was determined to be 58°C, by temperature gradient (56-62°C). In consequence, it was possible to run duplexed assays as described above, involving one Gram-Positive and one Gram-negative species each (*E. coli* + *S. aureus* and *K. pneumoniae* + *Enterococcus* spp.). These duplexed assays showed low replication variability and very low limit of detection, being able to detect DNA solutions with concentrations of 1pg/μL and in some cases 0.1pg/μL as we briefly described in Merino *et al* ^6^.

The ddPCR assay employed to detect and quantify *E. coli* and *S. aureus* did not amplify any of the other non-*E. coli* Enterobacteriaceae or any other species from the genus *Staphylococcus* other than *S. aureus*. The same was true for the duplex assay for *K. pneumoniae* and *Enterococcus* spp. In this case the ddPCR assay did not amplify other species from the genus *Klebsiella* or other non-*K. pneumoniae* Enterobacteriaceae and *Streptococcus agalactiae* (Table 1).

The spiked blood experiments showed that there is an almost perfect correlation (0.997 ≤ r ≤ 1.000, p≤0.001) between the number of CFU/mL of the bacterial culture and the number of gene copies/mL detected by ddPCR in each dilution tested (Figure 1). For almost all bacterial species tested ddPCR presented gene copies/mL in the same order of magnitude as bacterial culture results (CFU/mL). The results also indicate that the ddPCR assays used in this study have a limit of detection of 1-10 CFU/mL. In this study we determined that time from sample preparation to results was approximately 3.5h to 4h, significantly reducing the time to get actionable information (bacterial identification) compared to the gold standard blood culture technique (24h-48h).

**Figure 1.**
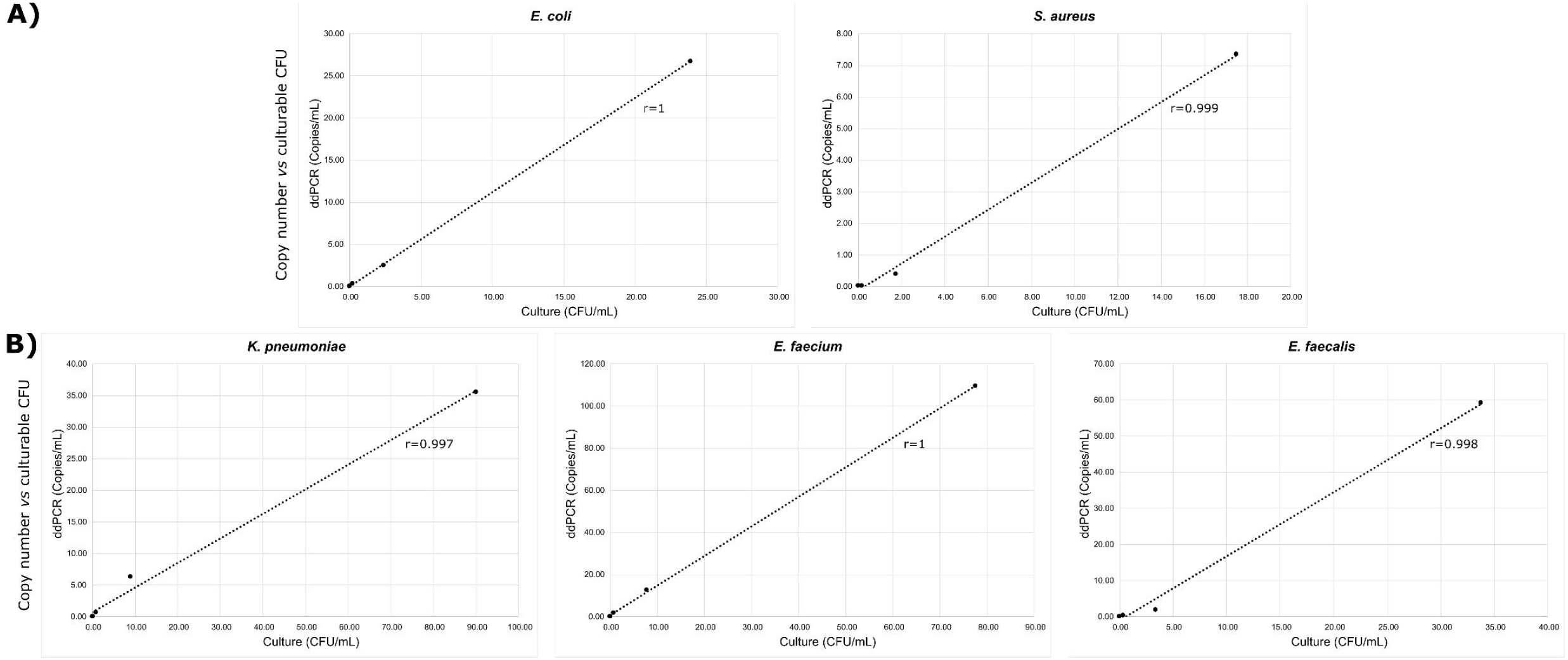
Correlation between the number of CFU/mL of the bacterial culture and the number of gene copies/mL detected by ddPCR. A) ddPCR assay to detect and quantify *E. coli* and *S. aureus*. B) ddPCR assay to detect and quantify *K. pneumoniae* and *Enterococcus* ^s^pp.

## Discussion

ddPCR demonstrated a very high sensitivity and specificity in this study. Regarding sensitivity, ddPCR assays showed a detection limit of 1-10 CFU/mL, similar to the few studies that have used other ddPCR assays ^2,13,14^ to detect DNA from bacterial pathogens in blood. This high sensitivity, make ddPCR assays ideal for detection and quantification of pathogens in sites where their initial load might be very small as BSIs and associated sepsis ^6,15^. Furthermore, this is the first study where a correlation has been performed between the number of CFU/mL obtained from spiked blood and copies gene/mL obtained from ddPCR. The almost perfect correlation we found between both quantification strategies indicates that we can approximate the number of bacteria present in patients’ blood.

Furthermore, the fact that the ddPCR assays tested in this study only amplified the intended bacterial species, indicates great specificity of these assays for each of the genera/species tested, confirming the results obtained in previous studies using the same primers/probes set ^10–12^. One of the main advantages of the described ddPCR assays over blood culture (gold-standard) diagnostic method is the time from sample to results we reported (3.5-4h *versus* 24-48h, respectively). The reported time of sample to results reported here is similar to that reported by other pilot studies using ddPCR assays to detect BSIs ^2,13^. The high specificity demonstrated by the ddPCR assays and the possibility to inform results in a timely manner to clinicians are important as it gives the analyst confidence to report the results obtained by this technique and in doing so helping clinicians to improve patients’ treatment.

Even though these assays were designed to detect the *E. coli, K. pneumoniae, S. aureus* and *Enterococcus* spp in blood they demonstrate the high performance of ddPCR, suggesting that this technology might be used for other applications (Figure 2), as: i) diagnostic of any type of infection ^6^, particularly those cases with low bacterial loads or difficult access to infection site (e.g. tuberculosis, meningitis or endocarditis), or in the case of sepsis where the positivity rate of current diagnostic techniques is low; ii) detection and quantification of antimicrobial genes or point mutations leading to antimicrobial resistance, allowing the reduction of the time from sample collection to results from at least 24-48h to a few hours, helping to guide and improve patients’ treatment ^2,13,14^: iii) severity stratification of the disease: high DNA loads have been associated to faster disease progression and greater mortality in patients with BSI ^9,16^; iv) distinguish between colonization and infection: quantifying bacterial DNA load might help to determine whether a certain opportunistic pathogen might be just colonizing a given site or if it is causing an infection instead (e.g. intestinal infections and lung infections); v) and finally, detection and quantification of the transcriptome of bacterial toxins: the detection and quantification of mRNA of *E. coli, Shigella* and *Clostridioides difficile* toxins might help with the diagnosis of infections by these bacterial species and also to assess disease severity, prognosis and guide treatment. Further studies specifically designed to test the performance of ddPCR to identify and quantify bacterial genomes directly from blood samples of patients with confirmed or suspected BSI are warranted.

**Figure 2.**
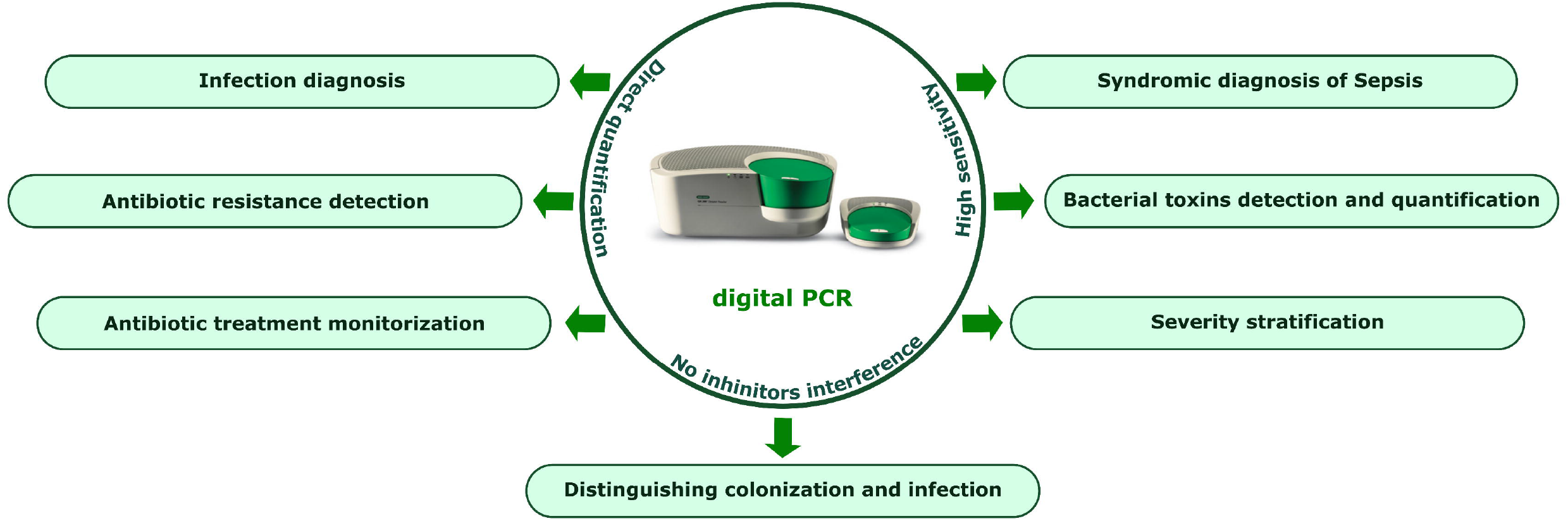
Possible applications of ddPCR for bacterial, viral and fungal infections diagnosis.

In this study we demonstrated that ddPCR assays to detect DNA from *E. coli, K. pneumoniae, S. aureus* and *Enterococcus* spp show a very low limit of detection, high sensitivity and specificity and can be used to identify and quantify DNA from these four bacterial genus/species directly from blood samples without the need of an intermediate step of bacterial culture. This pilot study reinforces the potential of ddPCR for the diagnosis and severity stratification of BSI and sepsis.

## Author contributions

**Ana P. Tedim:** Conceptualization; Methodology; Investigation; Formal analysis; Validation; Writing – Original draft; Funding acquisition. **Irene Merino:** Methodology; Investigation; Resources; Writing – review & editing. **Alicia Ortega:** Investigation; Resources; Project administration. **Marta Domínguez-Gil:** Resources. **José María Eiros:** Resources; Writing – review & editing. **Jesús F. Bermejo-Martín:** Conceptualization; Investigation; Formal analysis; Writing – review & editing; Supervision; Funding acquisition. All authors read and approved the final manuscript.

## Statements and Declarations

### Data availability statement

Data will be made available at a reasonable request.

## References

1. Timsit JF, Ruppé E, Barbier F, Tabah A, Bassetti M. Bloodstream infections in critically ill patients: an expert statement. Intensive Care Med 2020; 46: 266–84.

2. Wu J, Tang B, Qiu Y, et al. Clinical validation of a multiplex droplet digital PCR for diagnosing suspected bloodstream infections in ICU practice: a promising diagnostic tool. Crit Care 2022; 26: 243.

3. Martinez RM, Wolk DM. Bloodstream Infections. Microbiol Spectr 2016; 4.

4. Kalantar KL, Neyton L, Abdelghany M, et al. Integrated host-microbe plasma metagenomics for sepsis diagnosis in a prospective cohort of critically ill adults. Nat Microbiol 2022; 7: 1805–16. Available at: https://pubmed.ncbi.nlm.nih.gov/36266337/. Accessed January 10, 2023.

5. Kern W V, Rieg S. Burden of bacterial bloodstream infection-a brief update on epidemiology and significance of multidrug-resistant pathogens. Clin Microbiol Infect 2020; 26: 151–7.

6. Merino I, de la Fuente A, Domínguez-Gil M, Eiros JM, Tedim AP, Bermejo-Martín JF. Digital PCR applications for the diagnosis and management of infection in critical care medicine. Crit Care 2022; 26: 63. Available at: http://www.ncbi.nlm.nih.gov/pubmed/35313934.

7. Khatib R, Riederer K, Saeed S, et al. Time to positivity in Staphylococcus aureus bacteremia: possible correlation with the source and outcome of infection. Clin Infect Dis 2005; 41: 594–8.

8. Kim J, Gregson DB, Ross T, Laupland KB. Time to blood culture positivity in Staphylococcus aureus bacteremia: association with 30-day mortality. J Infect 2010; 61: 197–204.

9. Ziegler I, Cajander S, Rasmussen G, Ennefors T, Mölling P, Strålin K. High nuc DNA load in whole blood is associated with sepsis, mortality and immune dysregulation in Staphylococcus aureus bacteraemia. Infect Dis (Lond) 2019; 51: 216–26.

10. Weiss D, Gawlik D, Hotzel H, et al. Fast, economic and simultaneous identification of clinically relevant Gram-negative species with multiplex real-time PCR. Future Microbiol 2019; 14: 23–32.

11. Nijhuis RHT, van Maarseveen NM, van Hannen EJ, van Zwet AA, Mascini EM. A rapid and high-throughput screening approach for methicillin-resistant Staphylococcus aureus based on the combination of two different real-time PCR assays. J Clin Microbiol 2014; 52: 2861–7.

12. Cao Y, Raith MR, Griffith JF. Droplet digital PCR for simultaneous quantification of general and human-associated fecal indicators for water quality assessment. Water Res 2015; 70: 337–49.

13. Shin J, Shin S, Jung SH, et al. Duplex dPCR System for Rapid Identification of Gram-Negative Pathogens in the Blood of Patients with Bloodstream Infection: A Culture-Independent Approach. J Microbiol Biotechnol 2021; 31: 1481–9.

14. Zheng Y, Jin J, Shao Z, et al. Development and clinical validation of a droplet digital PCR assay for detecting Acinetobacter baumannii and Klebsiella pneumoniae in patients with suspected bloodstream infections. Microbiologyopen 2021; 10.

15. Wang M, Yang J, Gai Z, et al. Comparison between digital PCR and real-time PCR in detection of Salmonella typhimurium in milk. Int J Food Microbiol 2018; 266: 251–6.

16. Ziegler I, Lindström S, Källgren M, Strålin K, Mölling P. 16S rDNA droplet digital PCR for monitoring bacterial DNAemia in bloodstream infections. PLoS One 2019; 14.

